# Rapid evolution slows extinctions in food webs

**DOI:** 10.1101/107763

**Authors:** Peter C. Zee, Casey P. terHorst, Sebastian J. Schreiber

## Abstract

Historically, evolutionary changes have been thought to act on much longer time scales than ecological dynamics. However, a recent body of research has demonstrated that evolution that is rapid enough to dramatically affect ecological dynamics can lead to feedbacks between ecological and evolutionary processes. Thus, to understand the stability of ecological communities, we must also consider evolutionary change in the component species. Here, we use individual-based simulations of a quantitative genetic eco-evolutionary model to describe how trait evolution influences the stability of ecological communities. On short time scales, faster evolutionary rates decreased the probability of species extinctions as populations at low densities were rescued via trait evolution. However, on longer time scales, evolutionary had little effect on the number of extinctions. The extent of short-term evolutionary rescue depended on the source of trait variation; populations with variation generated through mutation experienced more rescue events and were less prone to extinction, relative to populations with only standing trait variation. Trait evolution leading to more rescued populations increased the stability of the community on timescales relevant to conservation. Our work highlights the importance of intraspecific trait variation and the evolutionary mechanisms maintaining this variation for community ecology, as well as management of declining populations in a community context.

## Introduction

While ecology and evolutionary biology each play central roles in our understanding of patterns of biological diversity, many ideas in these fields have developed independently. The time scales at which ecological processes and trait evolution operate have long been thought to be distinct, with ecological changes occurring over significantly shorter periods than evolutionary changes. However, recent work has made great strides in understanding how the conflation of these timescales affects both ecological and evolutionary dynamics. Rapid, contemporary evolution can affect ecological processes, such as population dynamics (e.g., Hairston et al. 2005; Turcotte et al. 2011), species interactions (e.g., Post and Palkovacs 2009; terHorst et al. 2010; Schreiber et al. 2011; Vasseur et al. 2011; Becks et al. 2012; Hiltunen et al. 2014; Patel & Schreiber 2015), community assembly (Pantel et al. 2015), and ecosystem function (Bassar et al. 2010; Miner et al. 2012; terHorst et al. 2014). In addition to our understanding of ecological processes, rapid evolution can have important conservation implications, as it may lead to evolutionary rescue, whereby a population is able to rebound from low population sizes following a change in the environment through changes in genotype frequencies (Lynch and Lande 1993; Gomulkiewicz and Holt 1995; Gonzalez et al. 2013; Carlson et al. 2014; Orr and Unckless 2014).

Rapid evolution and evolutionary rescue require intraspecific trait variation to provide the opportunity for evolution to occur within populations (Bolnick et al. 2011). In a closed population, the amount of variation subject to selection at any given time is a result of the standing genetic variation, with additional mutational input over time (Hermisson and Pennings 2005; Orr and Unckless 2014). If evolutionary change is dependent on waiting for rare mutations to arise, then it may be less likely to affect ecological outcomes over shorter time scales. In contrast, evolution can act on standing trait variation immediately. As such, many experimental eco-evolutionary studies have focused on trait evolution on standing variation (e.g., Hendry and Kinnison 1999; Reznick and Ghalambor 2001). Trait variation may also have ecological effects independent of evolutionary change, as more variable interactions can significantly alter the mean strength and outcomes of ecological interactions (Bolnick et al. 2011).

Consumer-resource interactions are a fundamentally important interaction in ecology, as they dictate energy flow through ecosystems (Begon et al. 2006; Estes et al. 2011). In natural communities, species are embedded in multispecies food webs of consumer-resource interactions (McCann 2011). These interactions can be complex and lead to both direct and indirect effects due to regularly observed phenomena such as omnivory (Pimm and Lawton 1978; Kratina et al. 2012) and intraguild predation (Polis et al. 1989; Arim and Marquet, 2004). The combination of these direct and indirect effects are important ecologically (Strauss 1991; Wootton 1994; Miller and terHorst 2012), but can also have important evolutionary consequences (Miller and Travis 1996; Walsh and Reznick 2008; Walsh 2013; Patel and Schreiber 2015; terHorst 2010; terHorst et al. 2015), resulting in complex evolutionary dynamics in multispecies communities.

How the complexity that arises from multiple species interactions affects the stability of those communities has been the subject of debate in ecology for decades (Pimm 1991; McCann 2000). The thought that diversity should stabilize communities, proposed by early ecologists (MacArthur 1955; Elton 1958) was challenged by May (1974), who demonstrated mathematically that increases in diversity result in a decreased likelihood of local stability in randomly interacting communities. Analogous stability criteria have since been described for a variety of more realistic ecological interaction structures (Allesina and Tang 2012). Ecologists have also investigated how trait variation and evolution affect ecological stability. Kondoh (2003) found that predators who behaviorally alter their foraging patterns to match an optimal niche increase species diversity and stabilize communities. This result echoes the early ideas that more diverse communities can be buffered through prey switching (MacArthur 1955; Elton 1958).

Here, we move beyond plastic changes in traits, such as behavioral modification, to investigate whether rapid trait evolution can have similar stabilizing effects in diverse ecological communities. We are particularly interested in whether rapid trait evolution can confer stability to communities through evolutionary rescue. While previous discussions of evolutionary rescue have focused on shifts in the abiotic environment (e.g., Bell 2013), we are interested in scenarios where populations can be evolutionarily rescued following changes in the community composition (Kovach-Orr and Fussman 2013; Yamamichi and Miner 2015). For example, prey that are driven to low densities by predators may recover because of ecological feedbacks that lead to predator-prey oscillations. Alternatively, prey populations may adapt in response to strong selection by predation pressure, and the surviving resistant genotypes increase the prey population size.

We built a stochastic, individual-based simulation model to study how the ecological and evolutionary effects of intraspecific trait variation influence stability in food web communities. We found that trait evolution stabilized these food web communities by slowing the rate of species extinctions on short time scales. On longer time scales, however, trait evolution had a less pronounced effect on the number of extinctions. A key mechanism underlying this increase in this transient increase persistence is population rescue via trait evolution. We did not find strong effects of species richness on the evolutionary effect on food web stability. Further, we found that this evolutionary stabilizing effect on communities relied predominantly upon the novel generation of genetic variation through mutation, rather than selection on preexisting standing variation.

## Models and Methods

### The Eco-evolutionary model

To study whether trait evolution confers stability to food web communities, we developed and simulated the dynamics of a stochastic, individual-based eco-evolutionary quantitative genetics model. The model consists of *N*_*B*_ basal species and *N*_*C*_ consumer species. Basal species are autotrophs (i.e., do not need to consume other individuals in order to reproduce), and are assumed to have no intraspecific trait variation, and thus do not evolve in our model. Consumer species, on the other hand, have intraspecific trait variation, and can evolve over time. These species are omnivorous (i.e., can potentially eat both basal and other consumer species), however, cannibalistic interactions are forbidden. Inclusion of cannibalism is beyond the scope of our current study, but we recognize it’s potential importance for ecological communities (Fox 1975; Rudolf 2007). While only consumer species are able to evolve in our model, both consumer and basal species exhibit ecological dynamics, and can go extinct.

The per-capita birth rate of the *i*-th basal species is *b(1-R*_*i*_*/S*) where *b* is the maximal per-capita birth rate, *R*_*i*_ is the density of the basal species, and *S* is the maximal number of occupiable sites available to species *i*. In the absence of predation, the per-capita death rates of all basal species are a constant *d*.

The attack rate of a consumer individual on a hetereospecific individual depends on the trait values of the interacting individuals (Nuismer and Doebeli 2004; Nuismer et al. 2005; Schreiber et al. 2011; Patel and Schreiber 2015). That is, an individual with trait value *y* attacks an individual with trait value *x* at the following rate *a(y, x)*:

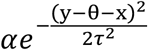

Where *α* is the maximum possible attack rate, *θ* is the optimal difference in trait values that maximizes the attack rate, and *τ* determines the degree of specialization of the predator (i.e., diet breadth) (fig. 1A).

**Figure 1:**
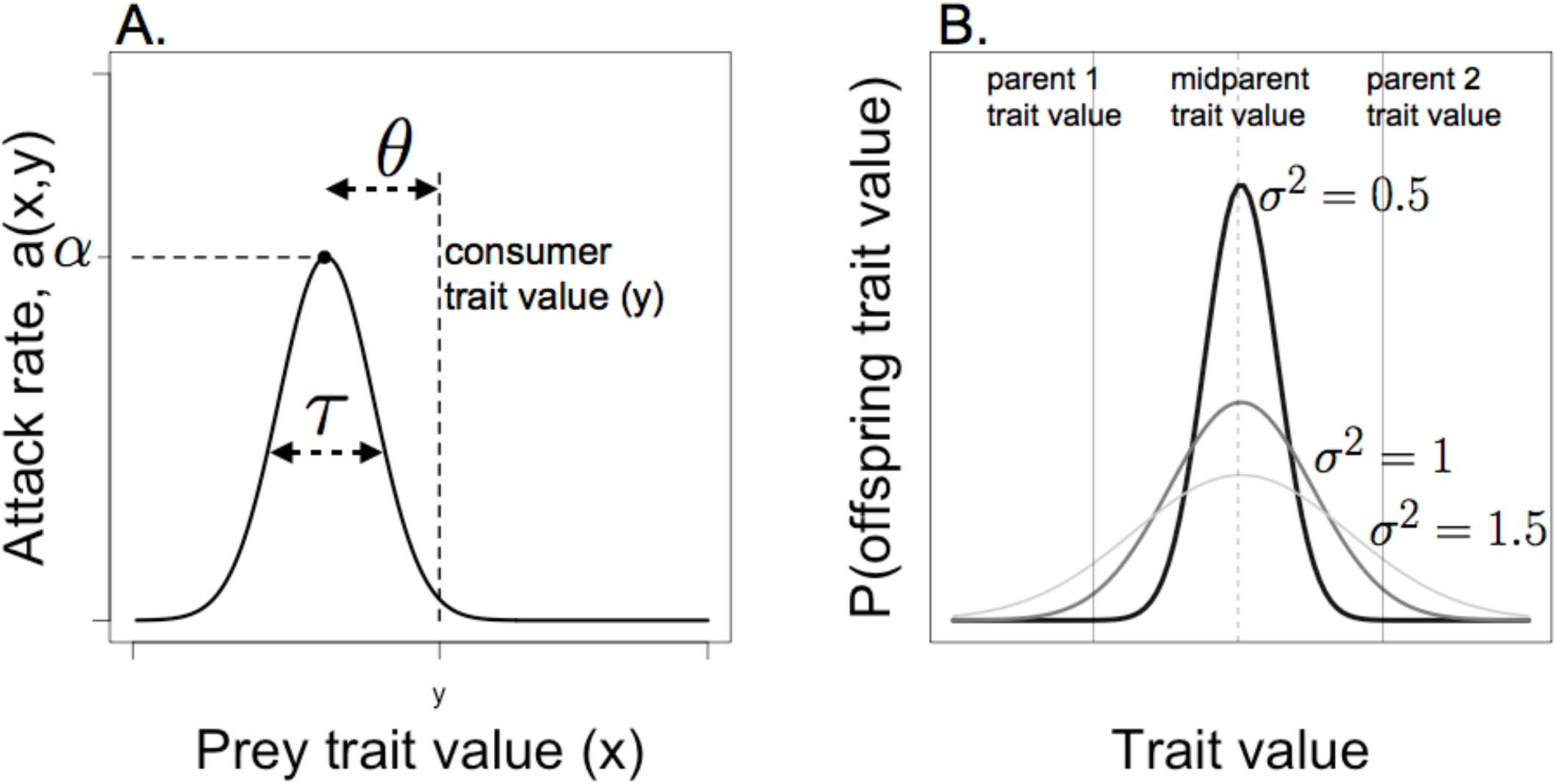
Attack rate and reproduction. (A) Attack rate function. (B) Trait values of consumer offspring are given by the deviation from the midparent mean trait value determined by the mutational variance *σ^2^*.

To describe the per-capita mortality rates due to consumption and the consumer birth rates, let *y*_*i*1_*,y*_*i*2_*,…, y*_*ici*_ be the trait values of *C_i_* individuals of consumer species *i*. Then the per-capita mortality rate due to consumption for an individual of consumer species *i* with trait *y* is

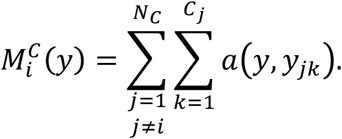

The per-capita mortality rate of an individual of basal species *i* is:

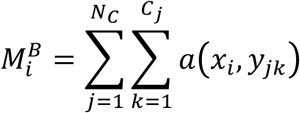

Where *x*_*i*_ is the trait value for all individuals of basal species *i*. The birth rate of an individual consumer of species *i* with trait *y* is proportional to its consumption rate:

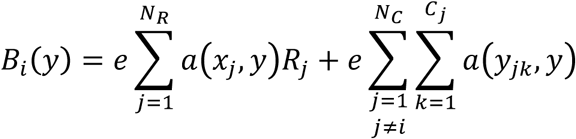

where the proportionality constant *e* corresponds to the conversion efficiency of a consumer. The first term of this sum corresponds to the consumption of basal species, while the second term corresponds to the consumption of hetereospecific consumers. When a consumer individual reproduces, it randomly mates with another conspecific individual. The trait value of the offspring is normally distributed around the mean of its parental trait values with mutational variance *σ*^*2*^ (Slatkin 1970; Lynch & Walsh 1998) (fig. 1B). With higher σ values, populations evolutionarily explore trait space more rapidly.

The model corresponds to a continuous-time Markov chain (Norris 1998) where individuals are updated at the previously described rates. That is, over a sufficiently small time interval Δ*t*, the probability of an individual of consumer species *i* with trait value *y* giving birth is (approximately) 
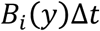
 and the probability of this individual dying is (approximately) 
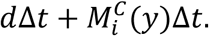
 The probability of an individual of basal species *i* giving birth is (approximately) 
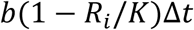
 while the probability of this individual dying is (approximately) 
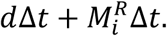
 The limit as Δ*t* goes to zero determines the continuous-time Markov chain. As doing exact simulations with Gillespie’s algorithm (Gillespie 1977) is computationally expensive, we use the tau-leaping approach (Cao et al. 2005, 2006) in which one performs simultaneous independent updates for sufficiently small Δ*t*. This method has been proven to approximate the distributional dynamics of the exact simulations (Anderson et al. 2011).

### Methods

We ran continuous-time, stochastic, individual-based simulations of communities following the model described above. In these simulations, we tracked changes in both population densities and mean trait values through time. We varied the amount of trait variation in the communities to assess the effect of rapid evolution in consumers. Finally, we ran alternative sets of simulations where trait variation is either generated solely through mutation or solely through pre-existing standing variation. In all cases, basal species trait values were fixed through time.

*Treatments and replication*. For both trait variation regimes, we simulated eco-evolutionary dynamics of 500 communities with *N*_*T*_ =20 (10 consumer and 10 basal species). In all cases, basal species had birth rates *b* = 0.25 and death rates *d* = 0.1, with population carrying capacities of 200 individuals. Consumer attack rate parameters were held constant for all consumer species: *α* = 0.001, *τ* = 0.1, and *θ* = 1. Consumer species converted consumed prey with efficiency *e* = 0.75 and died at rate *δ* = 0.01. Each simulation ran for 10,000 time steps. We scaled these time steps to the life expectancy for a typical individual of a basal species, as (*1/d*Δt), resulting in approximately 250 basal species generations per simulation. For each full set of *σ*^*2*^ values [0,1.5], the initial trait values for all species were independent draws from a uniform distribution on the interval [0,5]. We tracked ecological (changes in population densities) and evolutionary (changes in trait values) dynamics for each species in the community (fig. 2). While we primarily focused on communities with total initial richness of *N*_*T*_ = 20, we also simulated communities with total species richness ranging over *N*_*T*_ = {4, 6, 10, 20} (all with equal numbers of basal *N*_*B*_ and consumer species *N*_*C*_) and *σ*^*2*^ in [0,1.5], with 50 replicates for each treatment combination. Each community had initial trait values for the species (both basal and consumer) drawn randomly from a uniform distribution on the interval [0,5]. We found little effect of species richness on extinction probability, and no significant interaction with the effects of evolution. These additional results are presented in Appendix A.

**Figure 2:**
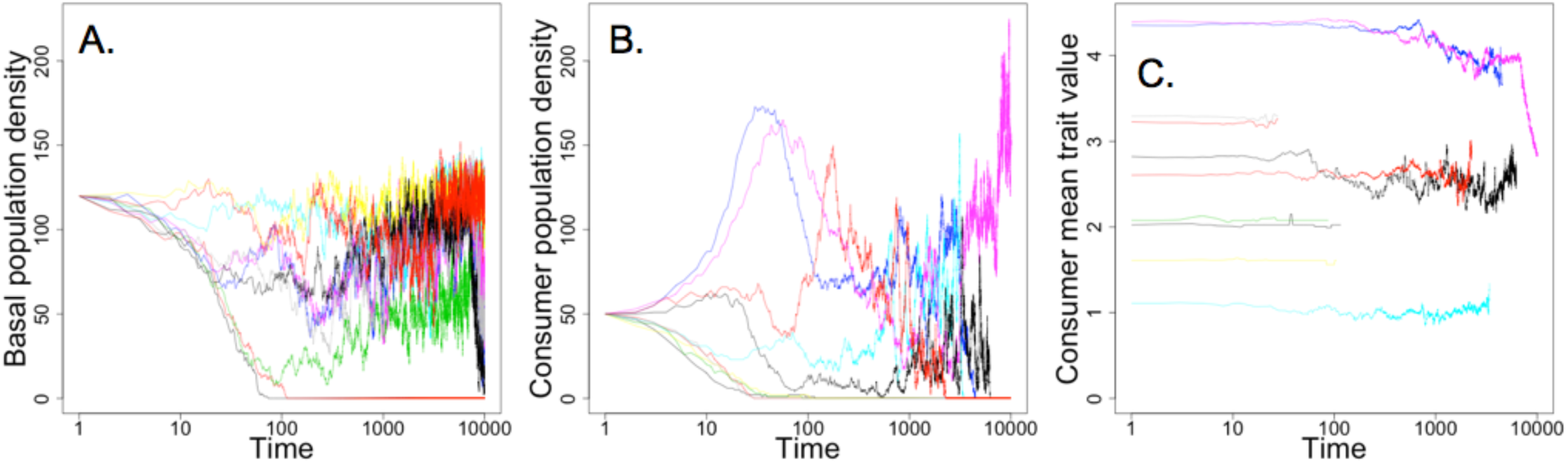
Representative ecological and evolutionary dynamics. (A) Ecological dynamics of basal species. (B) Ecological dynamics of consumer species. (C) Consumer trait evolution dynamics. Each curve shows dynamics for one consumer species. For the simulation shown, *σ^2^*=0.2. Note that time (x-axis) is on a log-scale

*Trait variation regime*s. To explore the effect of trait variation *σ*^*2*^ on community dynamics, we imposed two distinct regimes that determined how trait variation was generated. In the ‘standing variation’ regime, we introduced standing trait variation *σ*^*2*^ at the beginning of each simulation, and no subsequent trait variation was generated through mutation (i.e., the trait of all offspring is the midpoint of their parental traits). In the ‘mutation only’ regime, we initiated simulations with no initial trait variation, but mutations contributed to trait variation through time. In this ‘mutation only’ regime, all intraspecific trait variation arises via mutation over the course of the simulations. Comparison between these trait variation regimes allows us to determine whether lineage sorting of standing variation or continual mutation and selection contributes more to evolutionary effects on community dynamics (Appendix B).

*Extinctions and species persistence*. For both consumer and basal species, we measured stability through species persistence, i.e., the fraction of species persisting throughout the simulations. We measured both the rate of species losses as well as the total number of extinctions at the end of the simulations. The simulation data allowed us not only to determine the number of species extinctions in each community (extent of extinction), but also the timing of these extinction events (dynamics and rate of extinctions). We then compared how trait evolution affects the extent and rate of species extinction by looking at extinction patterns as a function of the mutational variance or initial standing variation (*σ*^*2*^). We defined the extent of extinction as the proportion of species at the beginning of the simulation that go extinct by the end of the simulation. To quantify the rate of extinction, we calculated the accumulation of extinction events through time and the average time for a community to experience 50% of its eventual extinction events.

*Population rescues*. To quantify the effect of trait evolution on consumer population rescues, we counted the number of times a population crossed a threshold density *L* (fig. 3). Each of these categories (i.e., number of threshold crossings) has a distinct biological interpretation. If a population never crossed the threshold, it means that the population persisted for the entire simulation while never falling to low densities. (‘Low’ is defined by the threshold density). If a population crossed the threshold density only once, we found that in our simulations the population always went extinct by the end of our simulations. Two crossings of the threshold density indicated that the population fell to low density, but was subsequently rescued and continued to persist to the end of the simulation. This scenario captures the iconic U-shaped population trajectory of evolutionary rescue (Gomulkiewicz and Holt 1995). Three crossings indicated a population was rescued, but then suffered subsequent extinction. More crossings (> 3) indicated multiple rescues from repeated low density, either with eventual persistence (an even number of crossings) or extinction (an odd numbers of crossings). We used a threshold density of *L* = 25 individuals, approximately 10% of typical peak population densities. We calculated the number of threshold crossings from smoothed population trajectory data to ignore small, random changes in population densities using the ‘filter’ function in R. While we quantified the number of threshold crossings, our presentation focuses on the number of rescues (the floor of half of the number of threshold crossings), the biologically more relevant quantity. For each rescue event, then, the population can either persist or subsequently go extinct.

**Figure 3:**
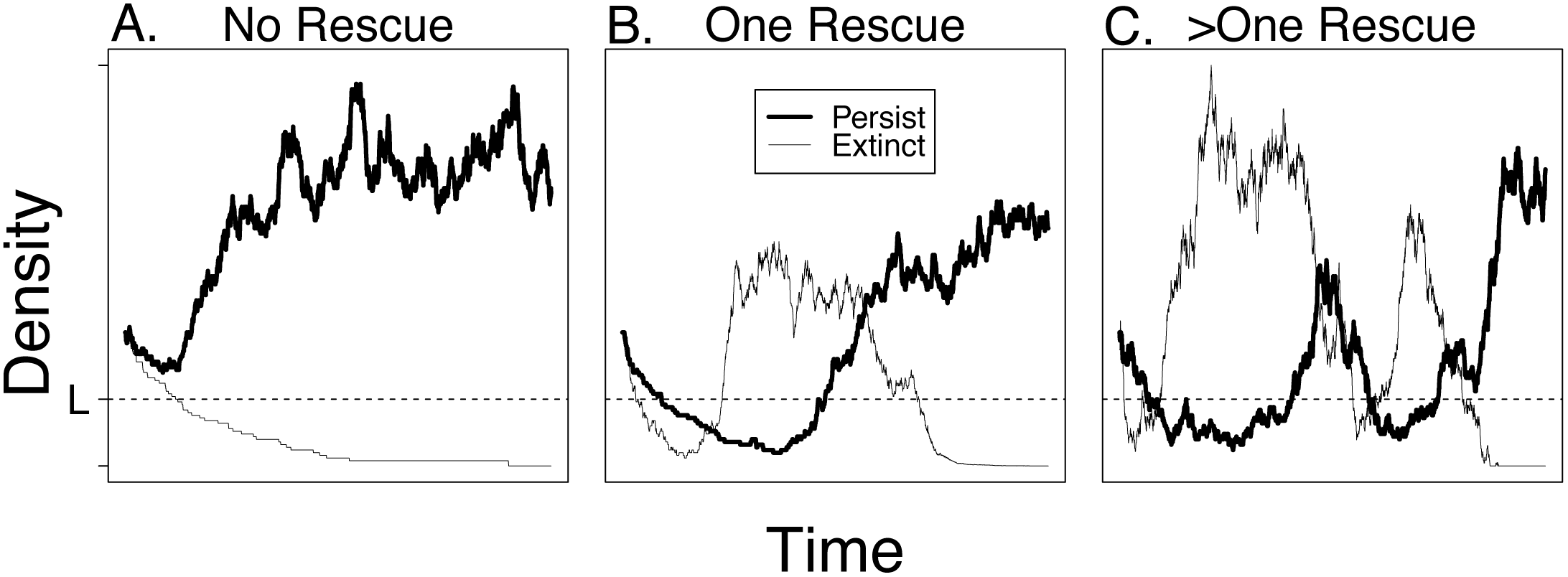
Classification of population rescues. Dashed line at *L* shows the low density threshold for population rescue. Each panel shows a classification of population rescues, with either eventual persistence (dark line) or extinction (light line). (A) No rescues. (B) One rescue. (C) More than one rescue.

*Dynamics of trait variation*. We examined the dynamics of trait variation with two methods. First, we quantified the standard deviation of population trait values for all populations through time for both the mutation only and standing variation regimes. This approach gave a general picture of differences in how trait variation was generated and maintained in the simulated communities. Second, we investigated whether the patterns of intraspecific trait variation correlated with population rescues. To do so, we measured trait variation in populations at the times at which they crossed the low density threshold *L*. That is, at the time when a population crossed *L*, we measured the standard deviation in its trait values. All simulations and subsequent data analyses were written and performed in R (R Core Development Team 2014).

## Results

### Extinction and persistence

The parameter controlling the rate of trait evolution (*σ*^*2*^) influenced the dynamics of population extinctions from communities by changing both the rate and extent of species losses (fig. 4). Increasing *σ*^*2*^ resulted in slowing of the consumer extinction rates (fig. 4B,E). We also tracked the basal extinctions over our simulation (fig. 4A,*D*). We found that at low *σ*^*2*^ values, basal extinctions are the highest, with no extinctions occurring at *σ*^*2*^ > 0.5. At low *σ*^*2*^ values (0.1 - 0.2), consumer populations drove basal species extinct, and had the potential to evolve to utilize new basal species, resulting in the continued accumulation of basal extinctions at these low levels of mutational variance (fig. 4B). This decrease in basal extinctions at higher values of *σ*^*2*^ can be attributed to the decreasing ability of a population to efficiently exploit the basal species when consumer populations have more variable trait values. As *σ*^*2*^ increases, the average attack rate of each consumer individual decreases (fig. C1). This decrease can be interpreted as a cost of trait variation (Schreiber et al. 2011).

**Figure 4:**
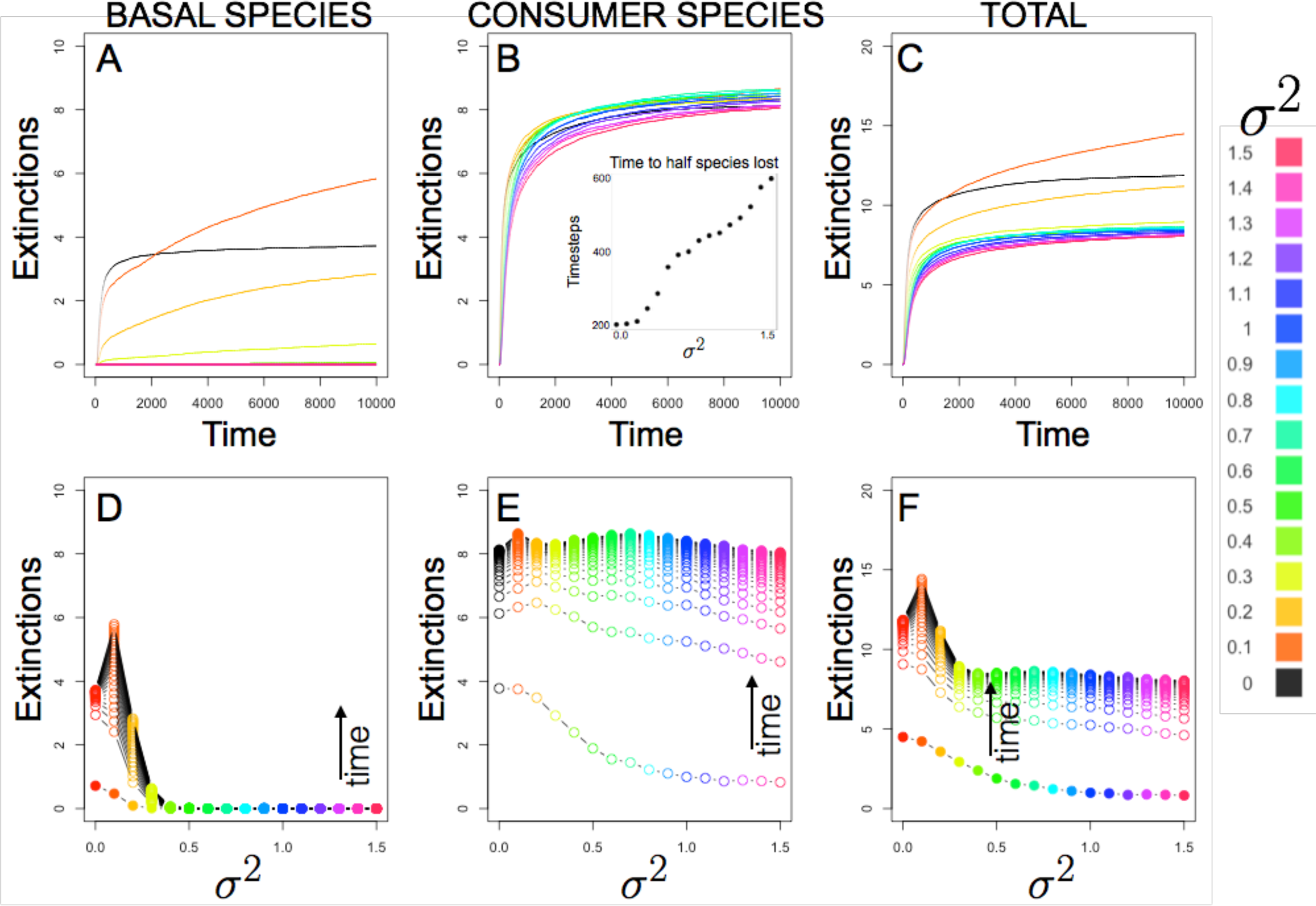
Extinction dynamics. (A-C) Each curve represents the accumulation of extinctions over time for different values of the mutational variance *σ^2^*. (D-F) The relationship between basal (B), consumer (C), or species (D) extinctions and *σ^2^* at different points in time. The uppermost curves in (B)-(D) show the mean number of extinctions at the end of our simulation runs. In all panels, colors represent different values of *σ^2^*.

As a result, we see that the number of basal species extinctions peaks at low values of *σ*^*2*^ (0.1), then quickly falls to zero as *σ*^*2*^ increases (fig. 4D). In contrast, the extent of consumer extinctions is relatively constant with respect to *σ*^*2*^ (fig. 4E). Figure 4D and 4E show the shifting relationship between *σ*^*2*^ and the number of extinctions over time. The bottom and top curves in each panel represent the extent of extinction at early and late time points, respectively in the simulations, across a range of *σ*^*2*^ values. For consumer species, we show that the effect of *σ*^*2*^ on the extent of extinctions becomes less negative over time (fig. 4E). However, for basal species extinctions continue to accumulate at lower values of *σ*^*2*^, leading to extinction peaks at these values of *σ*^*2*^ (fig. 4D). This peak at *σ*^*2*^ = 0.1 corresponds to the slight increase in the number of consumer extinctions, and is reflected in the dynamics of total community extinction (fig. 4F).

### Population rescues

For any given number of rescues, populations can either persist or eventually go extinct. We found that the number of populations that undergo rescue (2+ crossings) increases with *σ*^*2*^, while the number of populations that directly go extinct (one threshold crossing) or persist at high density without falling to low density (zero threshold crossings) decreases with *σ*^*2*^ (fig. 5A).

**Figure 5:**
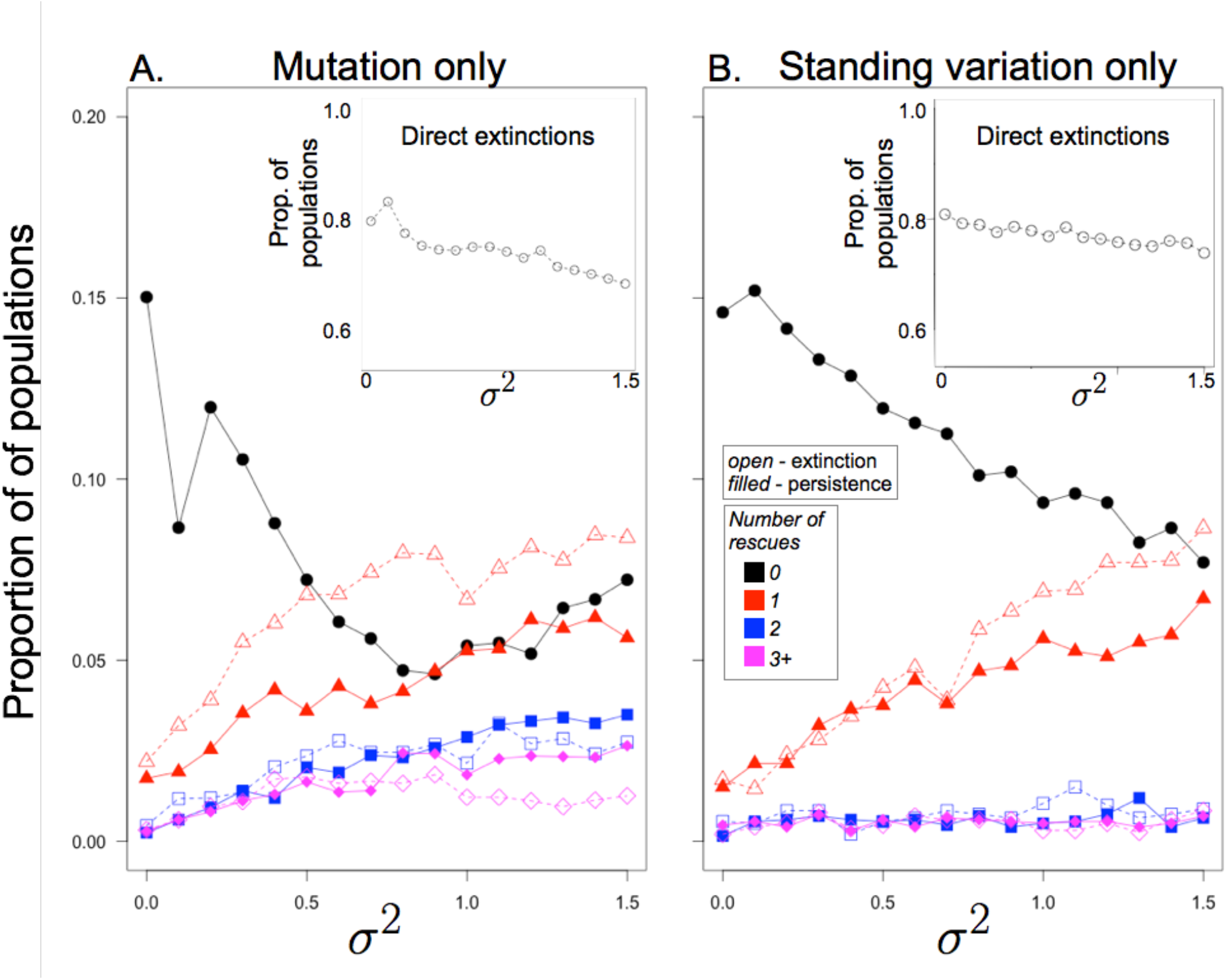
Evolution increases proportion of populations rescued. (A) Evolutionary effect on rescue under the ‘mutation only’ trait variation regime. Increased mutational variance (*σ^2^*) leads to more trajectories with rescues (and multiple rescues). (B) ‘Standing variation only’ trait variation regime. Here, probability of a trajectory having multiple rescues is not elevated. In both panels, the black points show a decrease in the populations that have zero crossing, i.e., persist at high density. In all cases, closed black circles represent the proportion. Upper panels (open black circles) in all cases show the proportion of populations that directly go extinct (one crossing). Note that the y-axis scales are different for lower and upper panels.

### Trait variation regimes

In simulations in which trait variation is generated via mutations, the proportion of populations that were rescued one or multiple times increased with *σ*^*2*^ (fig. 5A). By contrast, in simulations with only standing variation, the proportion of populations that were rescued once increased with *σ*^*2*^, but the proportion of multiple rescues (which is close to zero) was relatively unaffected by *σ*^*2*^ (fig. 5B).

Trait variation due to mutations or standing genetic variation differentially affected the proportion of populations that persisted without falling to low density (zero threshold crossings) (fig. 5). With mutational input, the proportion of these persisting populations decreased rapidly at low values of *σ*^*2*^, and then slightly increased at high values of *σ*^*2*^. With standing variation only, the proportion of these persisting populations decreased in a linear fashion with increasing *σ*^*2*^.

### Exhaustion of genetic variation

Mutations generated trait variation throughout the simulations as long as populations persisted, while trait variation was exhausted quickly when initial standing variation was the only source of trait variation (fig. 6A). Mutations generated trait variation to the same level of variation that existed at the beginning of the standing variation regime within 100 time steps (~2.5 basal species generations) at the beginning of the simulations. However, mutations continually renewed this variation, whereas without mutations, the variation was rapidly lost.

**Figure 6:**
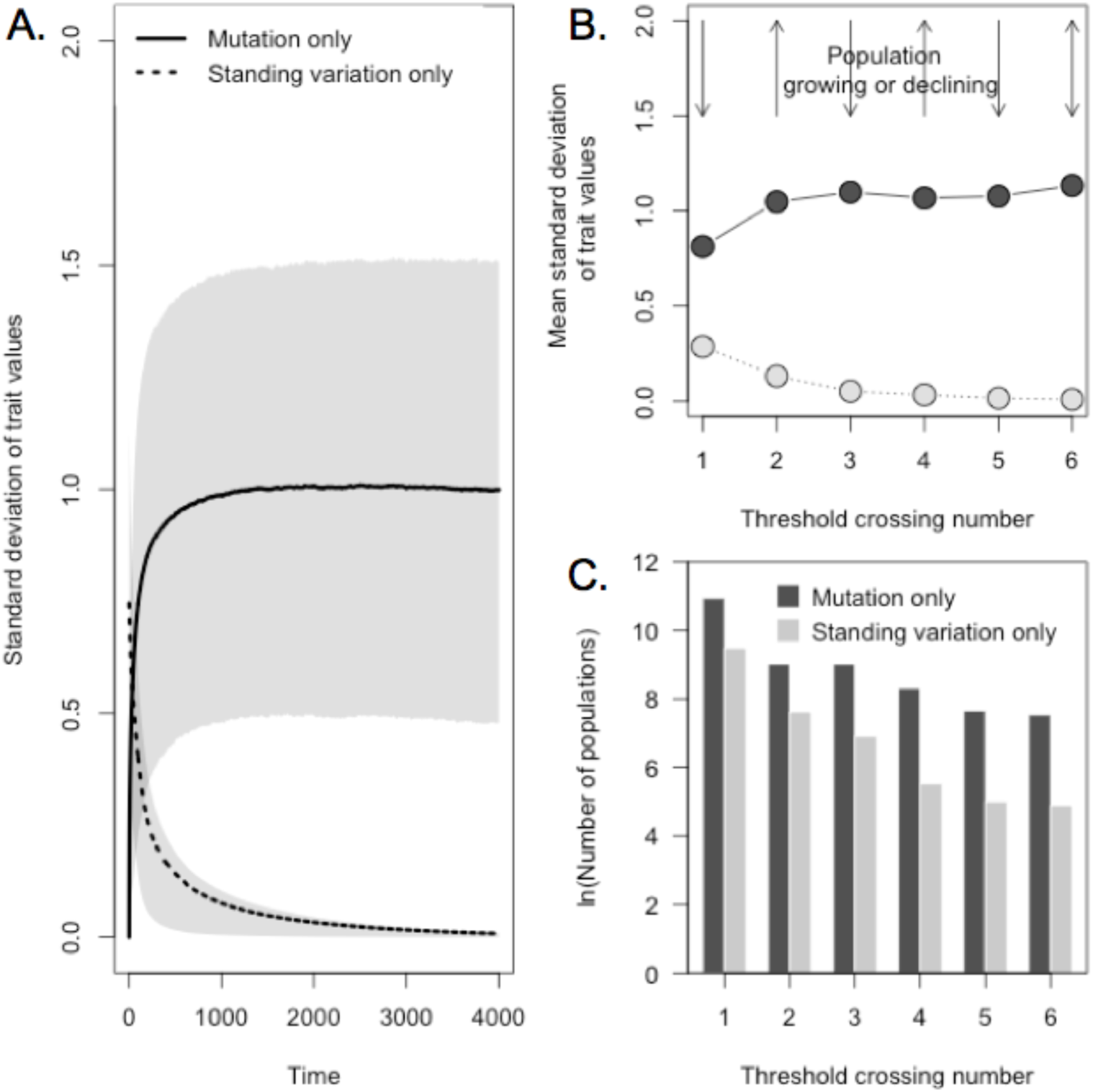
Standing variation only regime exhausts trait variation more quickly than mutation only regime. (A) The mean standard deviation of the trait values decreases to zero in the standing variation only regime, but stays level for the mutation only regime. Lower and upper bounds of shaded areas represent the first and third quartiles, respectively. (B) At the time of threshold crossings, there is significantly less intraspecific trait variation in the standing variation only regime (light) compared to the mutation regime (dark). Arrows at top represent whether the populations are increasing or decreasing in density at the time of threshold crossing. (C) Dark (mutation only) and light (standing variation only) bars represent the total number of populations for each regime with each crossing. Note the y-axis of (C) is on logarithmic scale.

At times of crossing the threshold density, the standard deviation of trait values was significantly higher for the mutation only regime (fig. 6B). In particular, at the time of secondary rescue events (4+ threshold crossings), the trait variation in the standing variation only regime fell to virtually zero (fig. 6C). By contrast, the standard deviations in trait values for the ‘mutation only’ regime were approximately equal for all secondary threshold crossings, with slightly lower values at the first crossing. This suggests that before the first threshold crossing, at which point the population is decreasing, mutations had not generated as much variation as later in the simulation. Indeed, with standing variation, the mean time to the first rescue event was significantly lower than the mutation only regime (Mann-Whitney U test, *p* < 0.0001; fig. B1).

## Discussion

Using individual-based, stochastic simulations of an eco-evolutionary model, we found that rapid trait evolution increases persistence by slowing the rate of species extinctions. The slower pace of extinctions found with increased mutational variance (i.e., evolutionary potential) is a result of evolutionary rescues from low population densities. While the rate of evolution – controlled by the mutational variance parameter *σ*^*2*^ – does not have a major effect on the number of eventual consumer extinctions in the long term, increased evolution substantially decreases the rate of extinctions on shorter time scales.

In the absence of evolution, extinctions occur very rapidly until the community reaches a stable state (fig. 4). If species in a community are unable to evolve, the initial community trait structure will determine which species go extinct. For instance, if the trait values of the initial community result in very low attack rates by a consumer species (no or few prey individuals available), then that population will starve and go extinct. Likewise, a consumer near the optimum on a basal species can rapidly drive the basal species extinct through consumption. This overexploitation can subsequently lead to the consumer species’ extinction.

At low values of *σ*^*2*^, consumers are able to evolve to overexploit basal species and drive them extinct. This overexploitation does not occur at higher mutational variances because the consumer attack rates are spread out among more basal species, allowing each population to maintain growth despite some consumption. Because there is no trait variation in basal species, increasing trait variation in consumer populations means that more consumer individuals are further away from the optimal trait difference from basal species (*θ*), thereby lowering average attack rates, and generating a fitness load on consumers (fig. C1). The consumers driving basal species extinct leads to an increase in subsequent consumer extinctions, resulting in a slight peak in the extent of consumer extinctions at *σ*^*2*^ = 0.1 (fig. 4E).

With trait variation, consumer species are able to evolve to consume new species if they drive one extinct. This can particularly be seen in basal species (fig. 4A), where the accumulation of extinctions reaches an approximate equilibrium quickly, while with trait variation, basal extinctions continue to accumulate. An analogous, but less pronounced, pattern is seen in the accumulation of consumer extensions (fig. 4B). However, as sigma values increase further, the load of trait variation suppresses the ability of consumers to drive prey species extinct (either basal or other consumers). This conclusion is further supported in the ‘standing variation only’ model, in which there is no mutational input, and therefore across all values of *σ*^*2*^, the curves for extinction accumulation level off more quickly (fig. B1). In the case of standing variation only, however, the load of trait variation vanishes because the variation is quickly lost (fig. 6).

Our results illustrate two ways that intraspecific trait variation can affect ecological communities. First, trait variation provides an opportunity for evolutionary change. We show that with inputs of trait variation through time, populations are able to avoid extinction through evolutionary change, leading to increased persistence. Second, beyond the opportunity for selection, trait variation can alter ecological interactions strengths, as exemplified by the reduced consumer attack rates at higher values of *σ*^*2*^ (fig. C1). Our simulations show how these two consequences of intraspecific trait variation can jointly affect community dynamics. This is most clearly demonstrated by the contrast in the number of basal species extinctions at low and high values of mutational variance (fig. 4A,D). These basal extinctions can then affect subsequent extinctions in the community. Populations with the same population trait means can have starkly different interaction strengths, reflecting the fitness load of increased *σ*^*2*^ (fig. C1; Schreiber et al. 2011). In nature, there is natural variation in many traits measured across taxonomic systems and environments (Lynch and Walsh 1998; Bolnick et al. 2011), and as such, ecologists should consider how trait variation affects species interactions within communities for more realistic predictions for outcomes of species interactions.

Introducing mutation and evolution into our simulations has different effects on species extinctions over short and long time scales. On shorter time scales, increasing *σ*^*2*^ stabilizes communities by slowing extinctions, but has diminishing effect over longer time scales. This difference among time scales of the effect of increasing *σ*^*2*^ is best illustrated in figure 4F, where the early negative relationship flattens over time. This suggests that early evolutionary and extinction dynamics result in a community structure that is relatively stable in the long term.

Extinction and evolution can alter the ecological connectedness in these communities, resulting in novel relationships among community members. In an individual-based population genetic model of predator-prey communities, Yamaguchi et al. (2011) also found that evolution could affect the number of extinctions in communities. In their two-trophic level system, they found that the effects of evolution on extinctions were dependent on the connectivity and genetic architecture. Evolution allowed consumers to change their prey use (i.e., gain or loss of trophic links), which could increase extinctions through altered competitive exclusion dynamics or decrease extinction via escape from competition among predators (Yamaguchi et al. 2011). We also found that evolutionary losses and gains of trophic links affected extinction dynamics, through overexploitation of prey species and evolutionary rescue of declining consumer populations. However, unlike their model, our communities were not strictly two trophic levels. In our model, because consumers can both evolve and eat each other, dynamics of trophic interaction losses and gains are complex. Interestingly, Yamaguchi et al. found that the specifics of the predator’s genetic system dictated whether the loss or gain of trophic links affected stability. Assessing the effects of variable genetic systems in our system is a compelling topic for future study (Schreiber et al. 2016).

### Mechanisms of rescue

Our method of quantifying rescue events as the number of threshold-crossings (fig. 4) does not provide direct information for the mechanisms underlying the rescue events. In the most intuitive rescue scenario, a focal population can be rescued from low density if the trait values of that focal species change such that some individuals are able to survive and reproduce (e.g., evolve to utilize a new prey resource). However, the mechanism of rescue may be indirect. For instance, a competing species could go extinct, or trait evolution in other community members could indirectly allow the focal species to rebound (e.g., Yamamichi and Miner 2015).

More simply, the multiple rescue events we found at higher *σ*^*2*^ values could arise from ecological dynamics in which predators abundances respond to prey abundances and vice versa, such as in predator-prey cycles. In such cases, the density of the ‘rescued’ consumer population would dip below the threshold density and recover as the basal species recover. However, the difference between the ‘mutation only’ and ‘standing variation’ regimes suggest that this is not the case. If multiple population rescues were driven by ecological dynamics, we would expect those dynamics to continue regardless of the regime of trait variation. In other words, if rescue events were driven purely by these ecological oscillations, we would not expect to see population rescues stop after the exhaustion of genetic variation in the standing variation regime (fig. 5). Comparison of the relationship between number of rescues and *σ*^***2***^ across the trait variation regimes supports that trait evolution is leading to the rescue events. This suggests that if populations are unable to continue to generate this intraspecific trait variation, they may soon face extinction risk, as they will not be able to be rescued from subsequent declines in density (fig. 6).

### Implications for conservation

We find that evolution can substantially slow the rate of extinctions on short time scales, but that it only modestly alters the extent of species loss from communities over longer periods of time. The short time scales over which we observe these strong effects are on the order of ~10 basal species generations, and thus relevant for most conservation efforts. The slowed extinction rate we found could provide critical time necessary for secondary ecological or environmental processes to act. In particular, this transient increase in persistence could be important when considering a broader landscape perspective (Thompson 2005). In this study, we have only simulated eco-evolutionary dynamics in a single focal community. We did not examine any effects of immigration or emigration among communities. When considering broader meta-community dynamics (Holyoak et al. 2005; Urban et al. 2008), the increased persistence though evolutionary rescue could have important effects on the maintenance of diversity at a larger scale. The slowed rate of extinction allows for more opportunity for rescue via immigration (Mouquet and Loreau 2002, 2003; Urban et al. 2008). Further, our model ignores other complexities that can alter eco-evolutionary outcomes, such as facilitative interactions (Bruno et al. 2003) and spatiotemporal environmental variation (e.g., Levins 1968; Mouquet and Loreau 2003).

Even with only standing genetic variation, we find that intraspecific trait variation is critical for evolutionary rescue of declining populations in the short term. In fact, evolutionary rescues occur significantly faster with standing variation (as opposed to waiting for relevant trait variation to accumulate through mutation) (fig. B2). This points to the maintenance of genetic variation in species of concern as an important aspect of conservation strategy (beyond the maintenance of variation for inbreeding avoidance). This suggests that in species with relatively slow generation times – and therefore reduced opportunity for rapid evolution on ecological timescales -- populations of these species with low variation may need input of variation if they are to be rescued through evolutionary mechanisms (Sgro et al. 2011; van Oppen et al. 2015; Whiteley et al. 2015).

### Future directions

Relaxing several assumptions of the current work opens compelling avenues for future research. We studied an abstract trait that governs ecological interactions among species in a food web community. Future work should seek to assign more explicit biological identity to the trait values. For instance, we could treat our trait values as body size, a variable trait that has been shown to determine feeding relationships across a wide breath of taxa and communities (Brose et al. 2006). However, with more explicit assignment of functional traits comes the attendant model complexities that would have to be addressed, such as trait dependent demographic parameters (i.e., rates of metabolism, birth, and death; Brose et al. 2006; Brose 2010).

Genetic rescue, as opposed to evolutionary rescue, can occur in declining populations by increasing fitness through the demographic contribution of immigrants. While such input of genetic variation can facilitate declining populations (van Oppen et al. 2015; Whiteley et al. 2015), this will depend on the genetic composition of the immigrants. In particular, when there is evolution in a focal/local community, if the immigrants are arriving from a source community where the selection pressures are not parallel to the local community, the effect of immigrants may have a variety of effects. It is possible that they could allow the population to recover (if the immigrating genetic variants facilitate response); alternatively, if the immigrants are phenotypically more ancestral, this could have the opposite effect of genetic rescue in which the immigrants constrain the evolutionary change, thereby increasing the likelihood of extinction. Further, immigration of basal species individuals could potentially stabilize consumer species that otherwise overexploit the basal species. Such studies of eco-evolutionary meta-community dynamics are compelling avenues for research (Urban et al. 2008; De Meester et al. 2016; Wittmann and Fukami 2016).

## Conclusion

We provide evidence that intraspecific trait variation and evolution significantly stabilize diverse ecological food web communities. Our eco-evolutionary simulations revealed that, on short time scales, the decreased extinction risk is a result of trait variation that allows species that have fallen to low density to be rescued via evolutionary changes. These results contribute to our understanding of how rapid evolution can generate patterns in ecological community dynamics that do not occur in the absence of intraspecific variation. Moreover, the slowing of species losses from diverse communities occurs on timescales relevant to conservation and management concerns, further stressing the value of evaluating concurrent ecological and evolutionary processes. Our results support the calls for accounting for intraspecific trait variation found in natural populations, and the resultant capacity for trait variation and rapid evolution to affect important ecological phenomena.

## Acknowledgements

We thank Swati Patel and other members of the terHorst and Schreiber research groups for helpful discussion. This work was funded by grants from the National Science Foundation to SJS and CPT (DMS-1312490), to SJS (DMS-1313418), and to CPT (OCE-1559105).

